# ACE2-Variants Indicate Potential SARS-CoV-2-Susceptibility in Animals: An Extensive Molecular Dynamics Study

**DOI:** 10.1101/2020.05.14.092767

**Authors:** Szymon Pach, Trung Ngoc Nguyen, Jakob Trimpert, Dusan Kunec, Nikolaus Osterrieder, Gerhard Wolber

**Author notes:** contributed equally (Szymon Pach, Trung Ngoc Nguyen). please send correspondence to Gerhard Wolber or Nikolaus Osterrieder.

## Abstract

Severe Acute Respiratory Syndrome Coronavirus 2 (SARS-CoV-2) emerged in late 2019 and since evolved into a global threat with nearly 4.4 million infected people and over 290,000 confirmed deaths worldwide.^1^ SARS-CoV-2 is an enveloped virus presenting spike (S) glycoproteins on its outer surface. Binding of S to host cell angiotensin converting enzyme 2 (ACE2) is thought to be critical for cellular entry. The host range of the virus extends far beyond humans and non-human primates. Natural and experimental infections have confirmed high susceptibility of cats, ferrets, and hamsters, whereas dogs, mice, rats, pigs, and chickens seem refractory to SARS-CoV-2 infection. To investigate the reason for the variable susceptibility observed in different species, we have developed molecular descriptors to efficiently analyze our dynamic simulation models of complexes between SARS-CoV-2 S and ACE2. Based on our analyses we predict that: (i) the red squirrel is likely susceptible to SARS-CoV-2 infection and (ii) specific mutations in ACE2 of dogs, rats, and mice render them susceptible to SARS-CoV-2 infection.

## Introduction

Severe Acute Respiratory Syndrome Coronavirus 2 (SARS-CoV-2) is responsible for the recent Coronavirus Disease (COVID-19) pandemic.^2^ SARS-CoV-2 and the related SARS-CoV (SARS-CoV-1), which caused the outbreak of severe acute respiratory syndrome (SARS) in 2002–2004, are different strains of the species *Severe acute respiratory syndrome-related coronavirus* of the family *Coronaviridae*. Coronaviruses (CoV) are enveloped viruses that present characteristic spike (S) glycoproteins on the surface of infectious virions that are essential for viral entry.^3^

S is a trimeric glycoprotein containing two functional subunits.^4^ The first subunit (S1) is responsible for binding to the host cell receptor, Angiotensin converting enzyme 2. The second subunit (S2) is responsible for the fusion of the viral and cellular membranes. The S1 subunit harbors the receptor-binding domain (RBD), which is responsible for binding to host cells.^3^ The SARS-CoV-2 RBD forms an antiparallel β-sheet structure, which is connected via short helices and loops.^5^ The contact to the host receptor is mainly established via loops known as the receptor-binding motif (RBM, Figure 1A). Compared to the SARS-CoV the RBM of SARS-CoV-2 appears to be less restricted in its conformation, because four of five prolines present in a loop structure essential for binding to the ACE2 protein are replaced by more flexible residues such as glycine in the S glycoprotein of SARS-CoV-2.^6^

**Figure 1.**
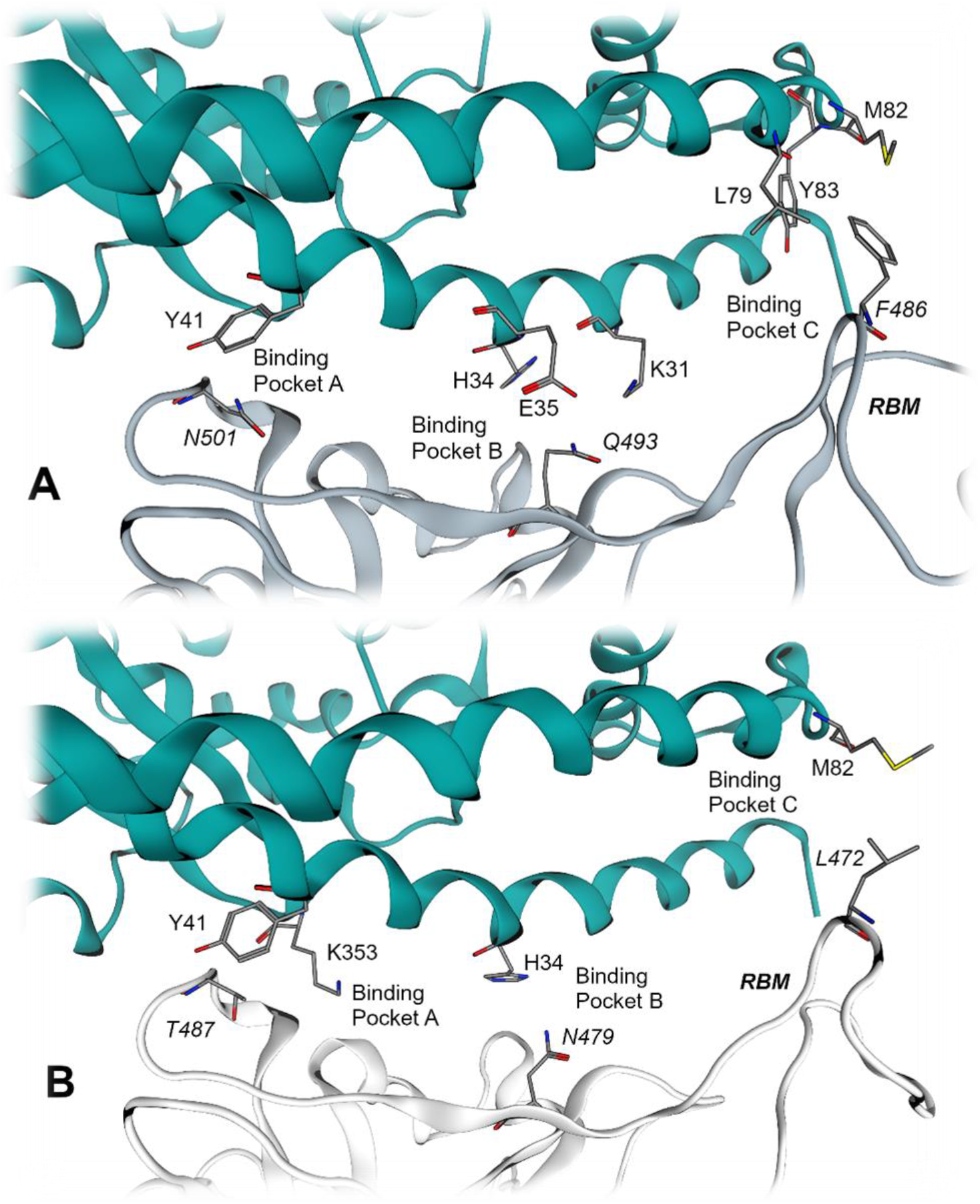
A: Three binding pockets (A, B and C) of the SARS-CoV-2 S-protein - human ACE2 binding interface; B: Analogous binding interface of SARS-CoV. Color-code: gray backbone-SARS-CoV, dark gray backbone-SARS-CoV-2, turquoise backbone-human ACE2. Residue names in italics belong to the viral RBD.

ACE2 is a membrane-bound zinc-metalloprotease expressed on the surface of cells not only in the respiratory tract but also in the heart, arteries, kidneys, and intestines were it acts as critical player in the endocrine renin–angiotensin system. Independent of its enzymatic function, ACE2 functions as the cellular entry receptor for coronavirus infections. The SARS-CoV-2 S shows nanomolar binding affinity to this cellular receptor.^7^ ACE2 forms a big substrate-binding cleft between the subdomains I and II.^8^ The SARS-CoV S binds to the ACE2 exploiting a non-competitive binding site on the outer site of subdomain I (Figure 1A, B).

Crystal structures of the RBD of the S in complex with human ACE2 have been described for both SARS-CoV ^9^ and SARS-CoV-2.^5^ The studies have shown that binding interfaces contain three homologous binding regions (pockets) in the ACE2-RBD interface, hereinafter referred to as binding pocket A, B, and C (Bp A, B, C). The RBD of the SARS-CoV S protein establishes hydrophobic contacts via its T487 in Bp A to a lipophilic pocket formed by K353 and Y41 of ACE2.^9^ Moreover, hydrogen bonds are formed between N479 and ACE2-residue H34 in Bp B, while lipophilic contacts are made between M82 (ACE2) and L472 in Bp C (Figure 1B).

The SARS-CoV-2 S features a binding interface consisting of three contact regions that are homologous to SARS-CoV. The binding interface is reported^5^ to utilize (i) hydrogen bonds in Bp A from N501 (homologous to T487 in SARS-CoV1) to Y41 of ACE2, (ii) a hydrogen bond network via Q493 (homologous to N479 in SARS-CoV) to ACE2-residues K31, H34, E35 in Bp B, and (iii) lipophilic contacts in Bp C via F486 (homologous to L472 in SARS-CoV) to ACE2-L79, M82, and Y83 (Figure 1A).

Recent reports confirm that in addition to primates, cats^10^, ferrets^10^, and hamsters^11^ are susceptible to COVID-19 infection, but dogs, pigs, chickens^10^, mice, and rats^11^ remain unaffected. Previous studies on SARS-CoV^9^ showed that differences in ACE2 sequences are responsible for the variable susceptibility observed in different species^9^. Based on these observations, we chose a structural approach to investigate how genetic diversity of ACE2 orthologs may affect the susceptibility of animal species to SARS-CoV-2 infection. On atomistic level, we have developed dynamic computational models for ACE2-RBD complexes of different species allowing us to anticipate the effects of amino acid sequence variation of ACE2 on viral entry.

## Results and Discussion

### Sequence comparison of animal ACE2 does not explain differences in COVID-19 susceptibility between investigated species

We focused on animal species based on (i) their importance as natural reservoirs for SARS-CoV-2 due to frequent contacts with humans and (ii) availability of studies reporting susceptibility to SARS-CoV-2 infection enabling discrimination between differences in our models. The multiple sequence alignment of canine, feline and human ACE2 orthologs revealed that the residues reported as hotspots^5, 9^ (referred to the human sequence positions 31, 34, 35, 41, 79, 82, and 83; canine positions are numbered one value lower than other species) show polymorphic mutations H34/33Y in dog and M82/81T in dog and cat. Since the difference in position 34/33 is present in dogs only, we searched for sequences harboring the same polymorphism. We found that ACE2 of common ferret (*Mustela pultorius*) also contains a tyrosine at this position. However, ferrets can be infected with SARS-CoV-2^10^, which suggests that the H34Y polymorphism does not prevent viral entry. Moreover, we compared ACE2 sequences from rodents (mouse, rat, hamster, and red squirrel) to sample additional binding pockets in the ACE2-RBD interface and predict susceptibility to SARS-CoV-2 of the red squirrel (*Sciurus vulgaris*).

Unfortunately, no protein structures are available for animal ACE2. To compare three-dimensional binding interfaces of animal ACE2-RBD complexes, we developed homology models of dog, cat, ferret, hamster, mouse, rat, and red squirrel proteins. We used the previously reported x-ray crystal structure of human ACE2 bound to the SARS-CoV-2 RBD (PDB-ID: 6M0J1^5^) as a template. All homology models show high quality with no major deviation in dihedral angles of the backbone (max. two Ramachandran outliers). Both outlier residues are located on flexible loops of ACE2 distal to the S binding site and represent polymorphic mutations from glycine in human crystal structure to serine in homology models. All homology models show comparable flexibility in molecular dynamics (MD) simulations (measured as root mean square deviation of backbone heavy atoms) to the human enzyme in the range of 3-5 Å.

Since the homology models of animal ACE2 are based on coordinates of human ACE2 in complex with the S RBD, we were able to superpose our models to the templates, yielding animal ACE2-SARS-CoV-2 RBD complexes. All binding interfaces of animal ACE2-S models remain in the same coordinate frame as the template crystal structure and thus comparable.

We compared ACE2 residues with direct contact to the RBD (distance of max. 4.5 Å) in all 3D complexes and searched for mutations possibly contributing to low SARS-CoV-2 susceptibility observed in dogs. However, polymorphisms were present in both non-susceptible and susceptible species, which did not explain the possibly lower affinity of RBD to dog, rat or mouse ACE2.

In the next step, we investigated dynamic properties of ACE2-RBD complexes. All structures were simulated in five MD simulation replicas (100 ns each) showing stable binding without dissociation events. For further analysis, we focused on three binding regions in the ACE2-RBD interface.

### Hot Spot Residue F486-Binding to Bp C Depends on ‘Depth’ and Conformation of Bp C

We discovered that the RBM shows considerably larger movements for canine and rat ACE2 complexed with the RBD when compared to other species. We focused on the hotspot residue F486 placed on top of RBM and occupying the lipophilic pocket (Bp C) of ACE2.

We chose the distance between the Cz-atom of F486 (position 4 on phenyl ring) and the Cb-atom of Bp C central residue 83 (or homologous position 82 in the dog model; Figure 2) as a surrogate parameter for F486 contacts. We observed three different states, which can be adopted by the F486-side chain: (i) ‘perfect’ fit into binding pocket (with distance of ca. 5-7 Å, ‘bound state’, Figure 2A), (ii) contact with ACE2 outside the binding pocket preferable with lipophilic or aromatic side chain of residue 78/79 (with distance of ca. 10-13 Å, ‘fixed state’, Figure 2B), and (iii) contact to lipophilic residues 27/28 and 78/79 with outwards rotated central Y82/83 in a ‘deformed state’ (with a distance of ca. 5-6 Å, Figure 2C). The contribution of the ‘fixed’ and ‘deformed state’ to RBD binding is yet unclear: Careful manual analysis of the performed MD simulations only revealed loose ACE2-RBD F486 contacts in these states, which suggests negligible binding contributions.

**Figure 2.**
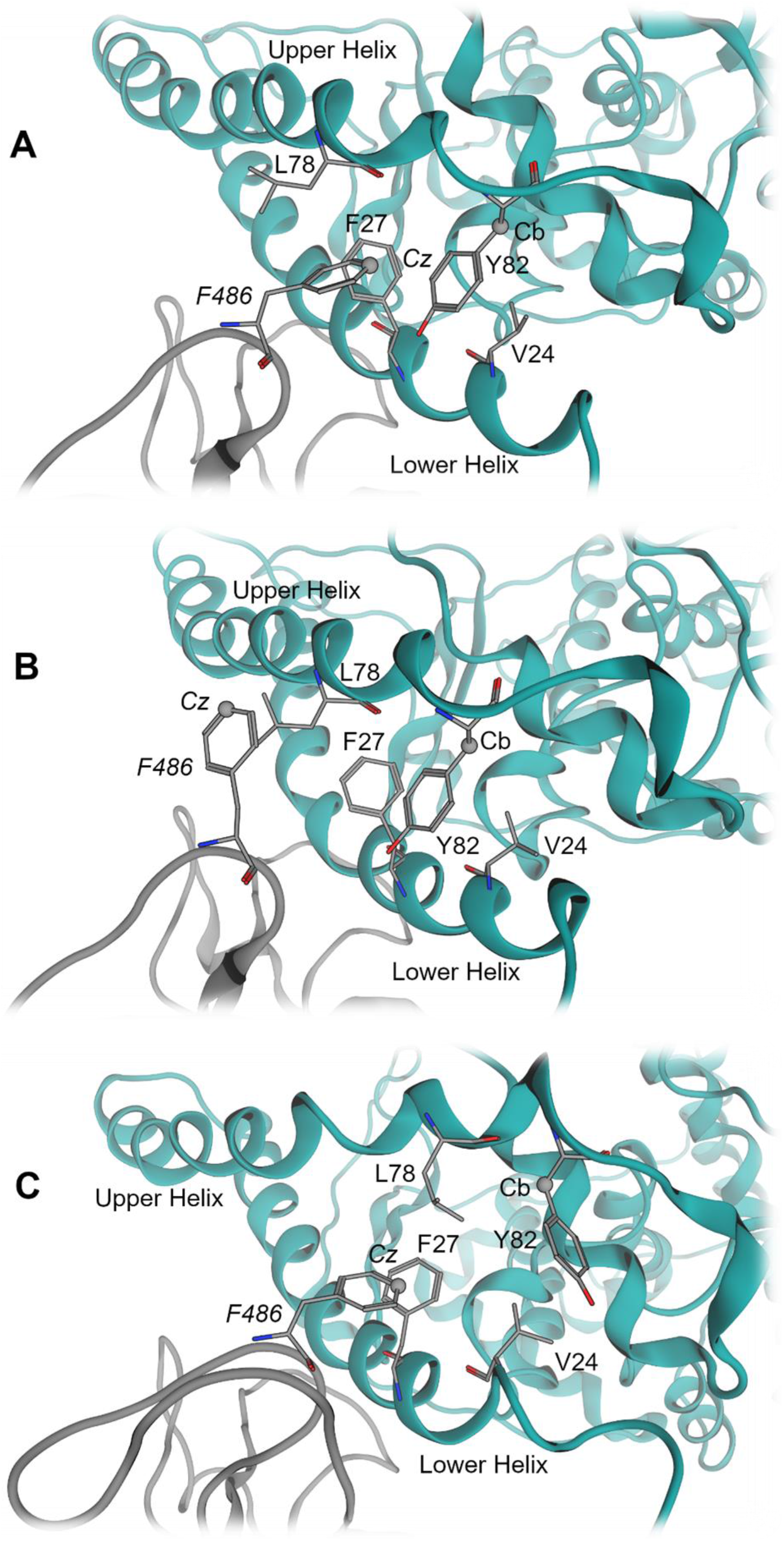
**A:** *Bound state* of F486 in the distance of 4-5 Å (measured between F486-Cz atom and Cb-atom of Y82); **B:** *Fixed state* of F486 outside the binding pocket; **C:** *Deformed state* of Bp C. The numbering of residues refers to the canine sequence.

From all analyzed species, cat, ferret, hamster, human, and mouse showed one or two peaks with a narrow distance distribution around 5-7 Å suggesting frequent occupation of Bp C (Figure 3). Although mouse ACE2 frequently occupies Bp C, we assume that weak interactions in the other two pockets (Bp A and B) might be responsible for the low SARS-CoV-2 susceptibility of mice. The simulations of rat ACE2-RBD complexes shows a dominant peak at around 11 Å implying F486 conformation in the fixed state. Canine ACE2-RBD complex simulations show three peaks suggesting transitions between bound, fixed, and deformed F486 states. All these results (with exception of murine simulations) correlate well with the susceptibility of the species to SARS-CoV-2.

**Figure 3.**
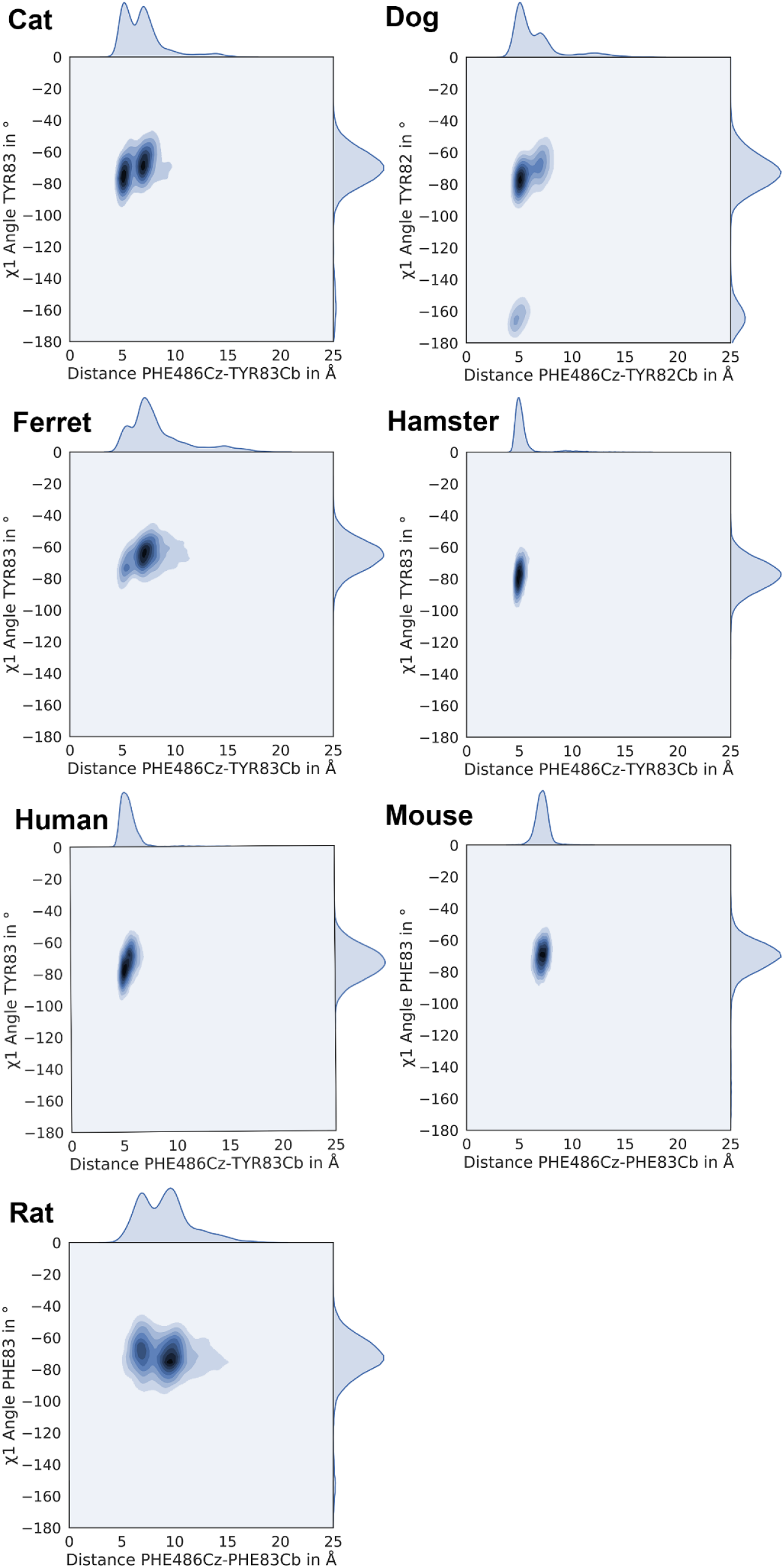
Occupation of binding pocket C indicated by kernel density plots (x-axis: distance F486 Cz - Y83/82 Cb) and rotamers of central residue 83 (or 82 in dog).

We surmised that the conformational changes of F486 in dog and rat simulations might be caused by either flattening or conformational deformation of Bp C. To validate this hypothesis, we calculated distances between the Cb atom of central residue 83 (or 82 in the dog ACE2) as the deepest point of Bp C and all side chain atoms of residues at positions 24, 79 and 82 (dog homologues 23, 78 and 81) flanking this pocket. We plotted the occurrence of the shortest distance per frame (SDpF) for each residue (Figure 4). From all three descriptors, SDpF between residues 83-79 (or 82-78 in dog) correlate well with Bp C occupation in rat ACE2. The residue 83-79 SDpF average of rat (7.1 Å) is the lowest distance in comparison with other species, which indicates a flatter Bp C. This would result in a suboptimal fit into the binding pocket and entails unbinding events of residue F486. MD simulations revealed that the polymorphism I79L (rat – all other species) might be responsible for the narrowing of rat Bp C. We observed that the methyl group of I79 restricts the Bp C and hinders fitting of F486.

**Figure 4.**
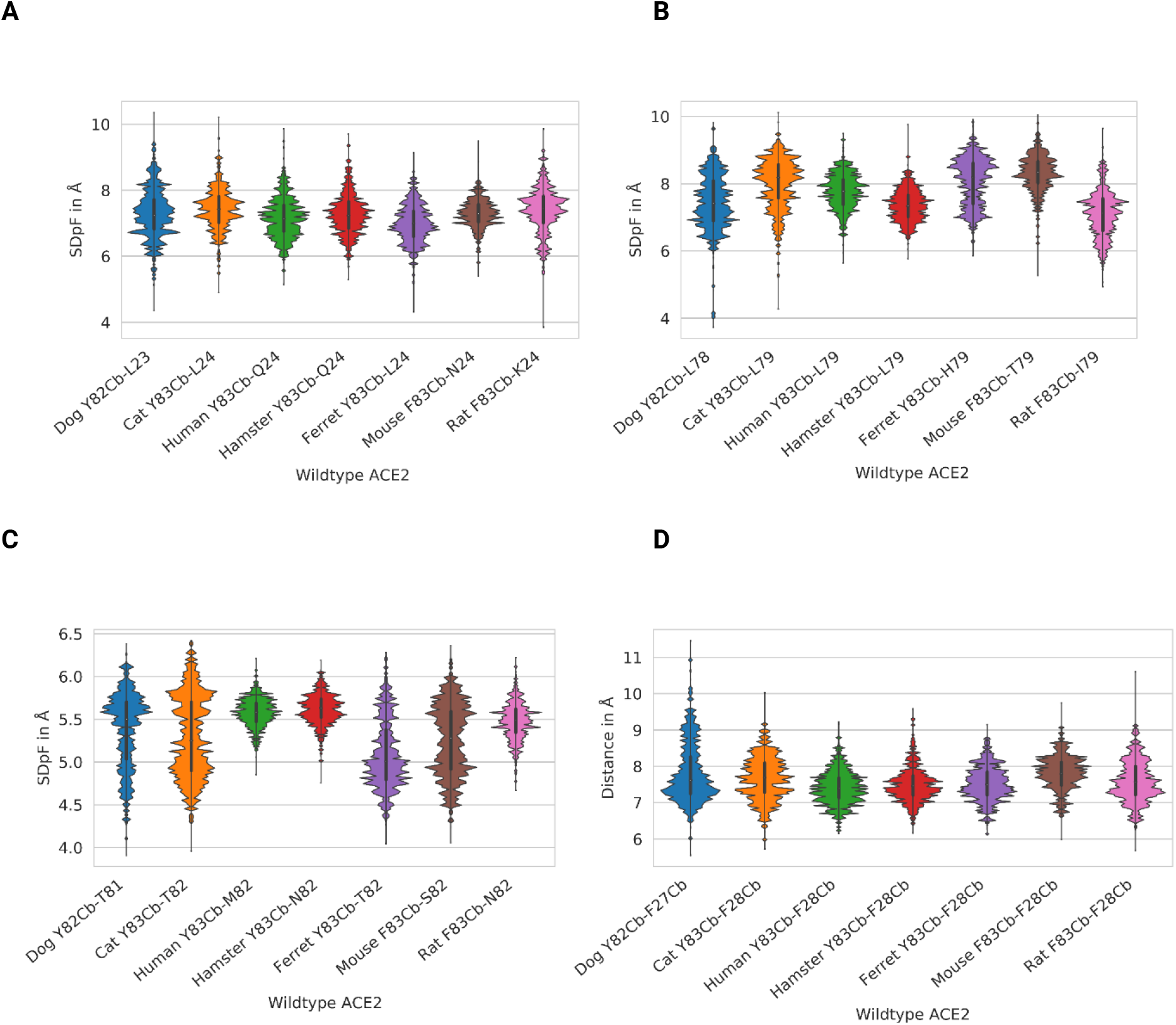
**A-C:** Distance distribution between Bp C key residues for analyzed wildtype ACE2 species. Shortest Distance per Frame (SDpF) between the Cb atom of residue 83 (or 82 in dog) and side chains flanking the binding pocket C (residues 24, 79, and 82 or 23, 78, and 81 in dog) representing the ‘depth’ of Bp C. **D:** Distance between the Cb atom of Y83 (or in dog 82) and the Cb atom of F28 (or 27 in dog) describing the deformation of Bp C.

The canine ACE2 shows a broad SDpF distribution between residue 82-78 with a minimal distance of 6 Å, which is comparable to rat simulations. However, this state is less frequent in canine simulations. Additional χ1 angle measurements of epitope 83/82 show that this residue rotates out of Bp C, which causes a deformation of the binding pocket in canine ACE2. We chose the distance between Cb atoms of residues 83 and 28 (or 82 and 27 in dog) localized in the upper and lower helix, respectively, forming Bp C (Figure 4, last panel). We surmised that the higher distance causes a larger shift between both helices and the resulting deformation of Bp C. In this state, the central residue Y83/82 can only establish weak interaction with F486. As expected, only canine ACE2-RBD simulations show higher distances than 9 Å for the Y83/82-F486 distance, which altogether suggests weak interactions. MD simulations revealed that the polymorphism V24/25A (dog – all other species) might be responsible for the outward rotation of canine Y82. The larger and more rigid V24 pushes Y82 out of Bp C and towards the N-terminus.

Subsequently, we analyzed the differences in distribution of residue 83-82 SDpF (or 82-81 SDpF in dog). Hamster, human, and rat show remarkably denser distance distributions (at approximately 5.5 Å) than other species. The sequence comparison shows that the three species express a long and non-branched side chain residue (asparagine in hamster and rat, methionine in human) at position 82 (dog 81). These amino acids might stabilize F486 in the ‘bound state’ with Bp C by steric effects. The most favorable residue at position 82/81 for binding F486 seems to be the methionine present in human ACE2, which also increases lipophilic contacts at this position.

### Mouse Mutation E/D30N and K31N in Bp B Disrupts Hydrogen Bond Network and Salt Bridges Found in Other Species

Due to contradictory findings in murine Bp C models, we surmise that the low SARS-CoV-2 susceptibility observed in mice might be caused by unfavorable or less frequent interactions in Bp B and/or A. Hence, we searched for polymorphisms exclusively occurring in the mouse ACE2 sequence. We found two mutations in murine Bp B (E/D30N and K31N), which are surrounded by charged amino acids. Both mutations replace a charged with a neutral residue, suggesting that RBD stabilizing salt bridges cannot be formed. Further analysis of MD trajectories of susceptible species led us to the hypothesis that (i) residue E/D30 can interact with K417 of RBD and (ii) K31 interacts with E35 within the same helix of ACE2 introducing stable amino acid pairs coordinated by viral residue Q493. Our hypothesis is supported by distance (SDpF) measurements between residues 30 (ACE2) - 417 (RBD) (Figure 5) and 31 (ACE2) - 35 (ACE2) / 31 (ACE2) - 493 (RBD) (Figure 6).

**Figure 5.**
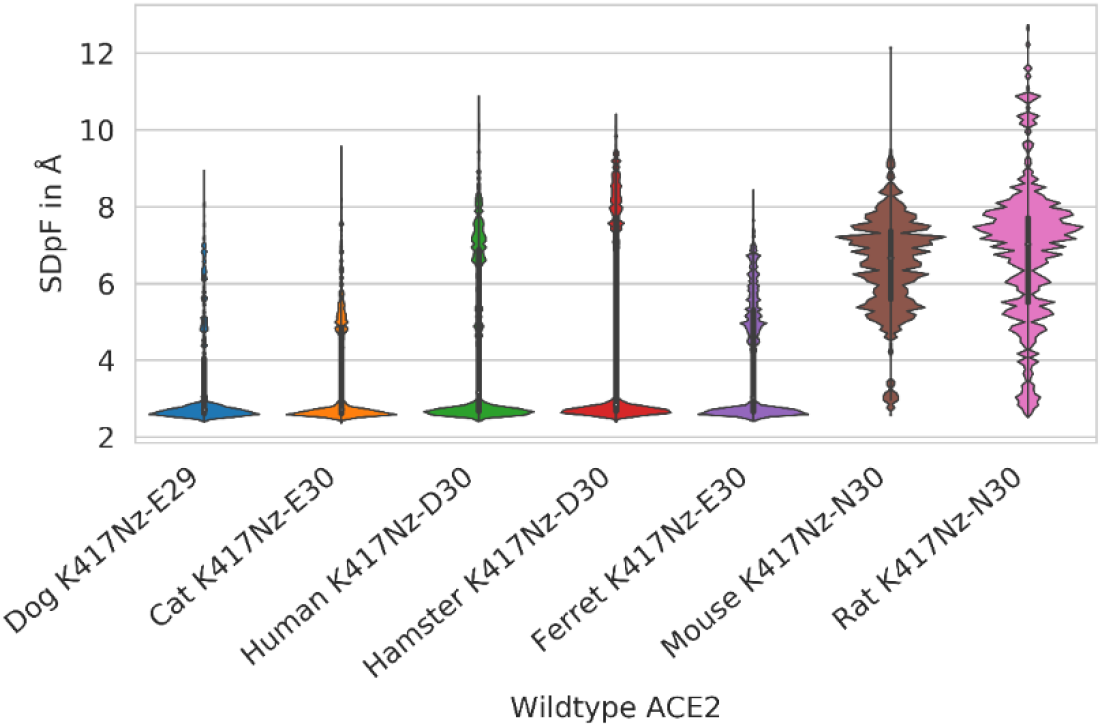
Shortest Distance per Frame (SDpF) between Nz atom of RBD K417 and side chain of ACE2 residue 30 (or 29 for dog) as surrogate parameter for interactions between these residues in the Binding pocket B for wild type ACE2.

**Figure 6.**
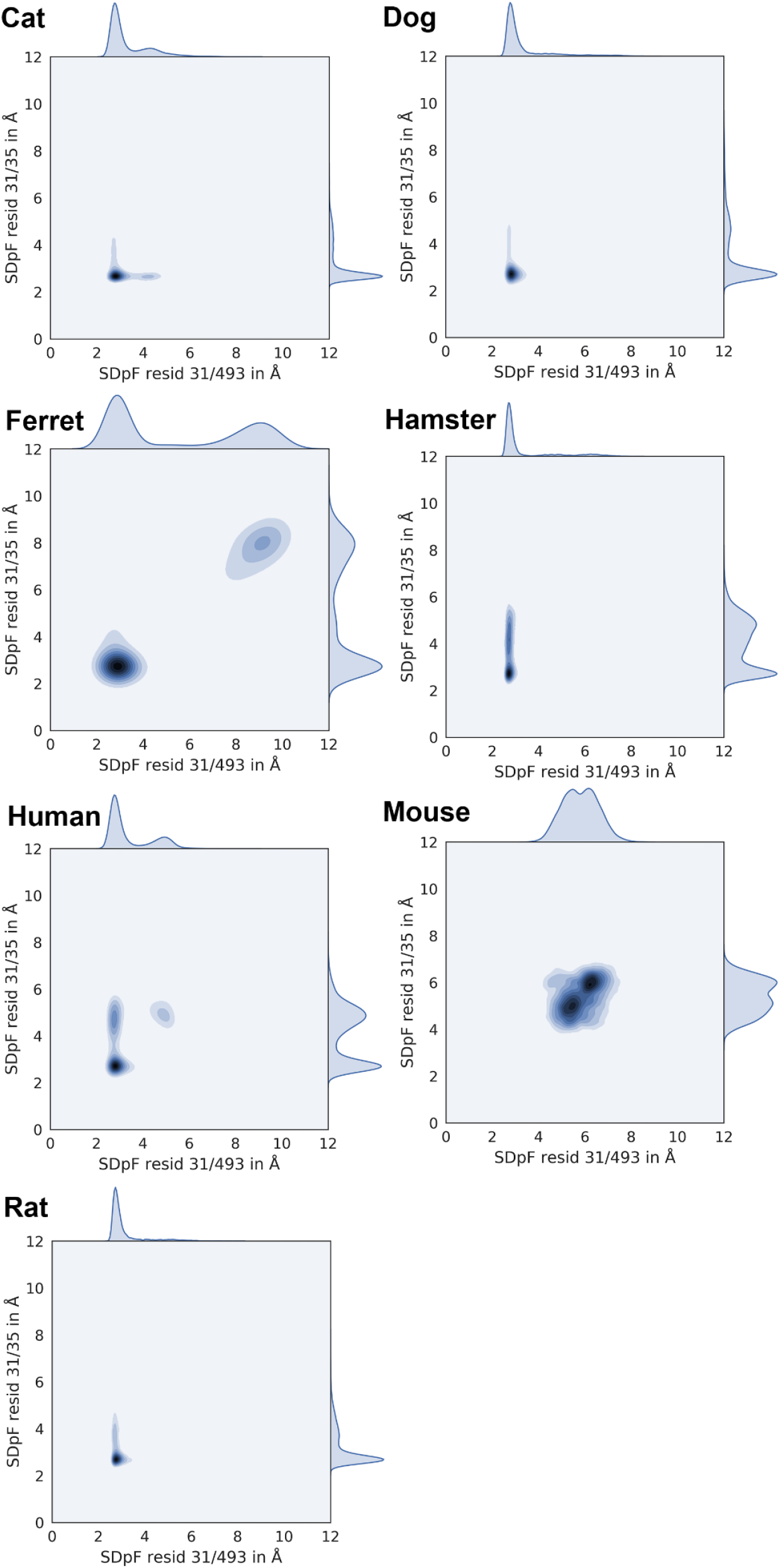
Interactions between ACE2 residue 31 and RBD residue Q493 summarized by kernel density plots (x-axis, SDpF: Shortest Distance per Frame between Nz K31 or Cg N31 - side chain of Q493) and interactions between ACE2 residues 31 and 35 (y-axis, surrogate parameter: distance Nz K31 or Cg N31 - side chain of E/D35).

Simulations of the murine protein complex show a considerably greater distance between residues N31, E35 and Q493 compared to other species (6 versus 3 Å). This indicates disruption of the hydrogen bond network in Bp B. Similarly, the mutation E/D30N occurring in mouse and rat breaks a salt bridge to K417 of RBD and the main distance peak occurs at 5-7 Å (in contrast to other species showing close contacts at ca. 3 Å). Due to larger distances for mouse and rat residues to RBD, water might invade the binding pocket resulting in lower S-protein RBD affinity to murine ACE2.

In addition, we found that a serine residue occurs at position 27 in rat ACE2 while more lipophilic threonine epitopes are found in other species. Since residue 27 is surrounded by lipophilic residues of the RBD (F456 and Y489), we surmise that the T27S polymorphism in rat might contribute to less favorable interactions in Bp B.

### Position 353 Regulates Hydrogen Bonding in Bp A and Explains Rat/Mouse Difference

To our knowledge, the role of residues involved in interactions within Bp A could not be clarified so far.^6^ We therefore compared the ACE2 sequence of rat and mouse with other species and identified the mutation K353H as relevant difference. Inspecting MD trajectories of rodent ACE2-RBD complexes, we observed that a lysine side chain in hamster ACE2 establishes a salt bridge to D38. Similar to the residue pair K31-E35 in the Bp B, which is coordinated by viral residue Q493, the pair K353-D38 interacts with RBD residue Q498. This region of Bp A is surrounded by a hydrogen bond network comprised of ACE2 residues 37, 41, 42, 355 and RBD residues 449, 496, 500, 501, 505. We assume that, according to the O-ring theory^12^, this hydrogen bond network ‘seals’ the hotspot K353/D38 (ACE2) - Q498 (RBD) and prevents water from invading the protein-protein interface. The histidine at position 353 in mouse and rat decreases the average number of hydrogen bonds established in the Bp A (7 in hamster versus 3 or 4 in rat or mouse, respectively; Figure 7).

**Figure 7.**
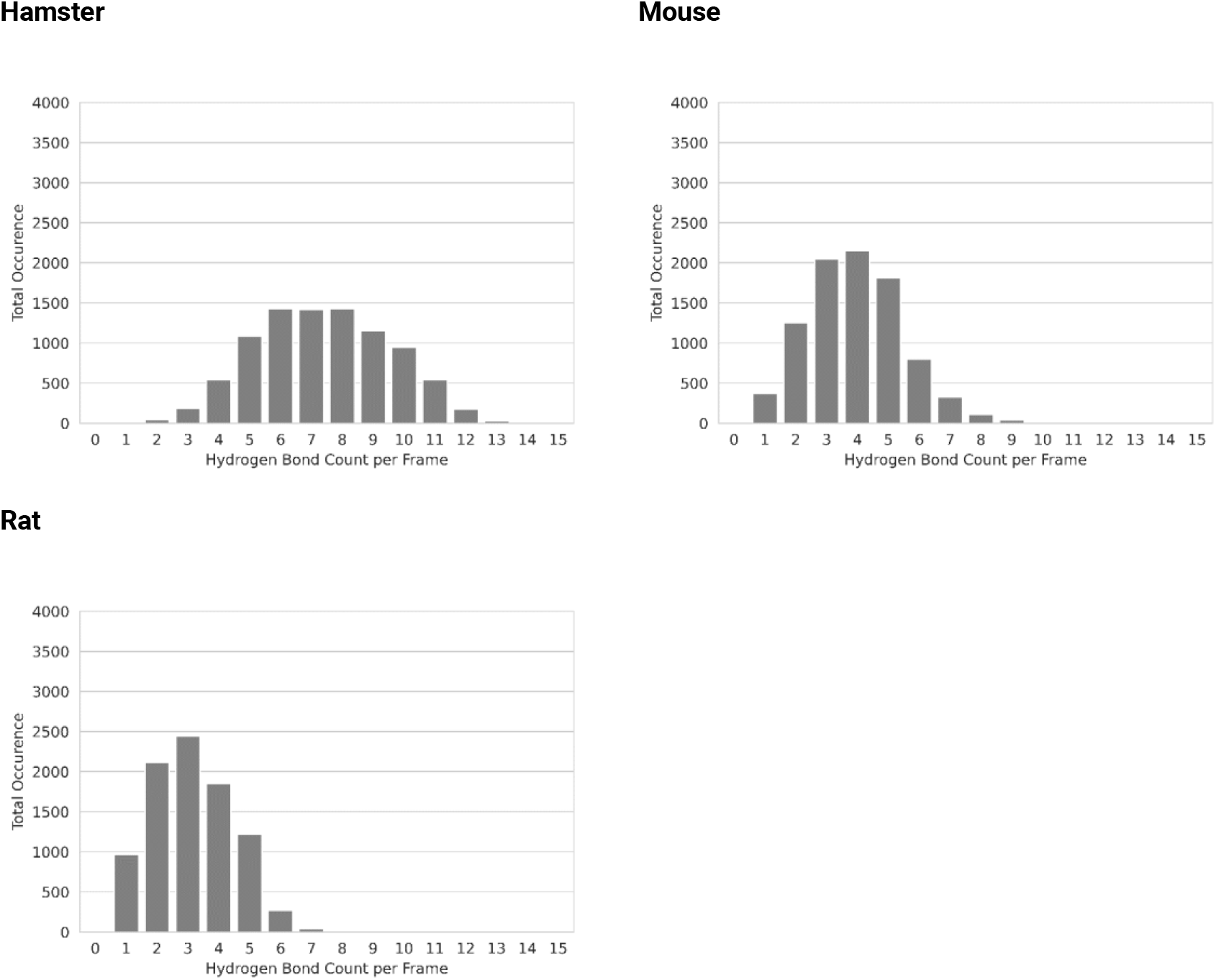
Total hydrogen bond count distribution in binding pocket A between ACE2 residues 37, 38, 41, 42, 353, 355 and RBD residues 449, 496, 498, 500, 501, 505 for hamster, mouse, and rat simulations.

Since histidine is less basic than a lysine, the formation of a salt bridge to D38 is less likely. MD simulations show that the histidine residue is too short to establish hydrogen bonds with D38 (ACE2) or Q498 (RBD). D38 can rotate out of the Bp A and water can reach the binding site, which results in less frequent interactions between the two proteins. This could explain why the RBD shows lower affinity to mouse and rat ACE2.

### Analysis of Squirrel ACE2 Suggests High Chance of Susceptibility to SARS-CoV-2

Red squirrels can be broadly found in urban environments in Europe. We therefore strived to predict squirrel ACE2 binding to S from SARS-CoV-2 since no information about susceptibility of the red squirrel to SARS-CoV-2 has been published yet. We prepared a red squirrel ACE2-RBD complex similarly to other species and conducted the workflow described above; (i) the depth parameters (83-79-SDpF) and occupancy of Bp C by RBD F486, (ii) distance plots between residues 30 (ACE2) - 417 (RBD) and 31 (ACE2) - 35 (ACE2) / 31 (ACE2) - 493 (RBD) in Bp B and (iii) H-Bond counts for Bp A show similar values to that registered in human ACE2-RBD complexes (Figure 8). Our analyses of Bp A-C indicate red squirrel is highly susceptible to infection with SARS-CoV-2.

**Figure 8.**
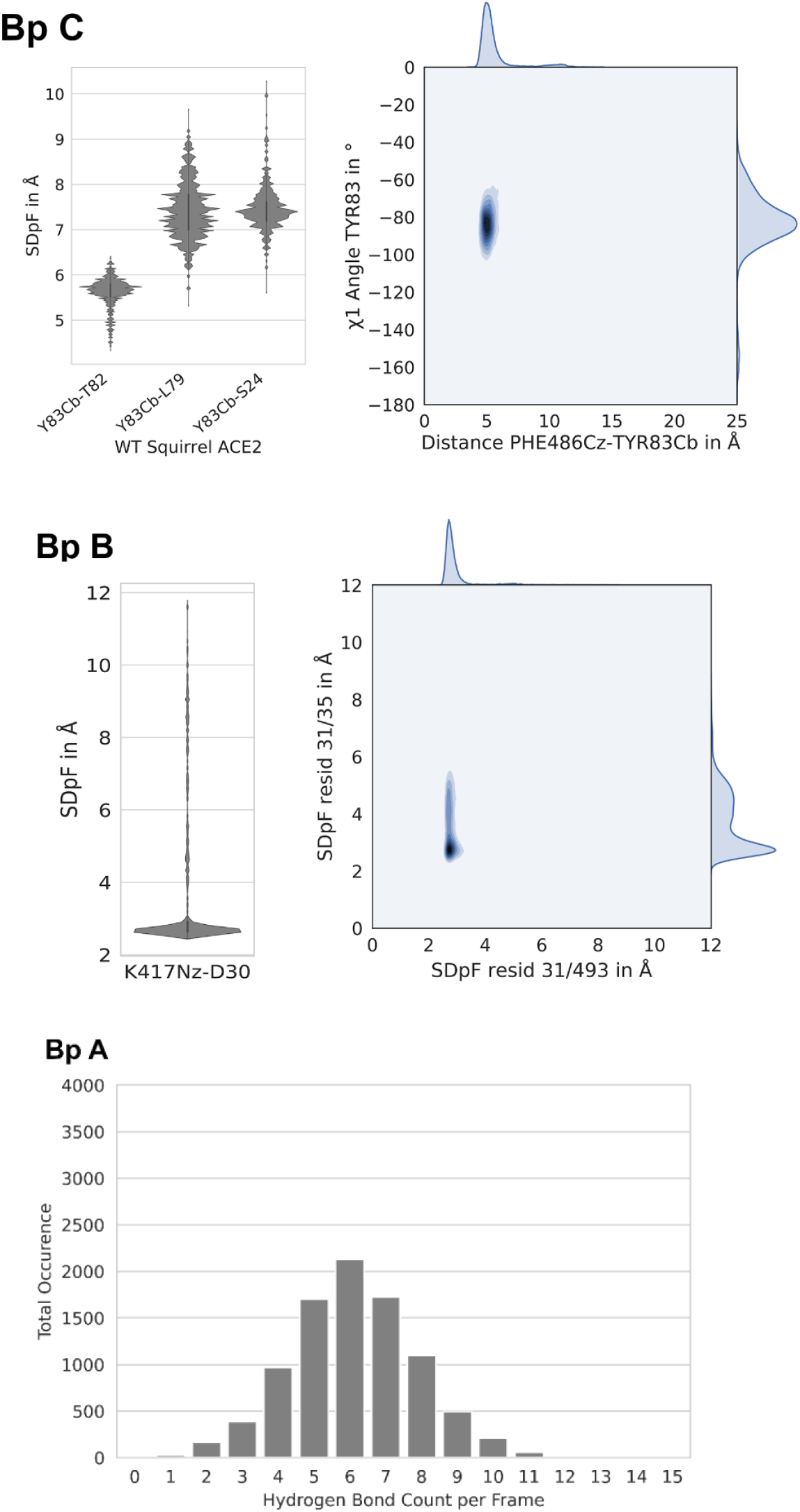
Descriptors calculated for red squirrel ACE2-RBD simulations. Bp C: Shortest Distance per Frame (SDpF) between Cb atom of Y83 and sidechains of residues flanking the binding pocket; Bp B: Distance distribution for interactions between Nz atom of RBD K417 and side chain of ACE2 residue 30, and kernel density plots summarizing the interactions between ACE2-residue 31 and RBD residue Q493; Bp A: histograms representing total occurrence of hydrogen bonds in binding pocket A between ACE2 residues 37, 38, 41, 42, 353, 355 and RBD residues 449, 496, 498, 500, 501, 505.

### Prediction of Gain-of-function Mutations

Based on our descriptors and sequence comparisons, we suggest mutations contributing to SARS-CoV-2 susceptibility in unaffected species (Table 1). We have investigated some of these mutants in-silico with dynamic models.

**Table 1.**
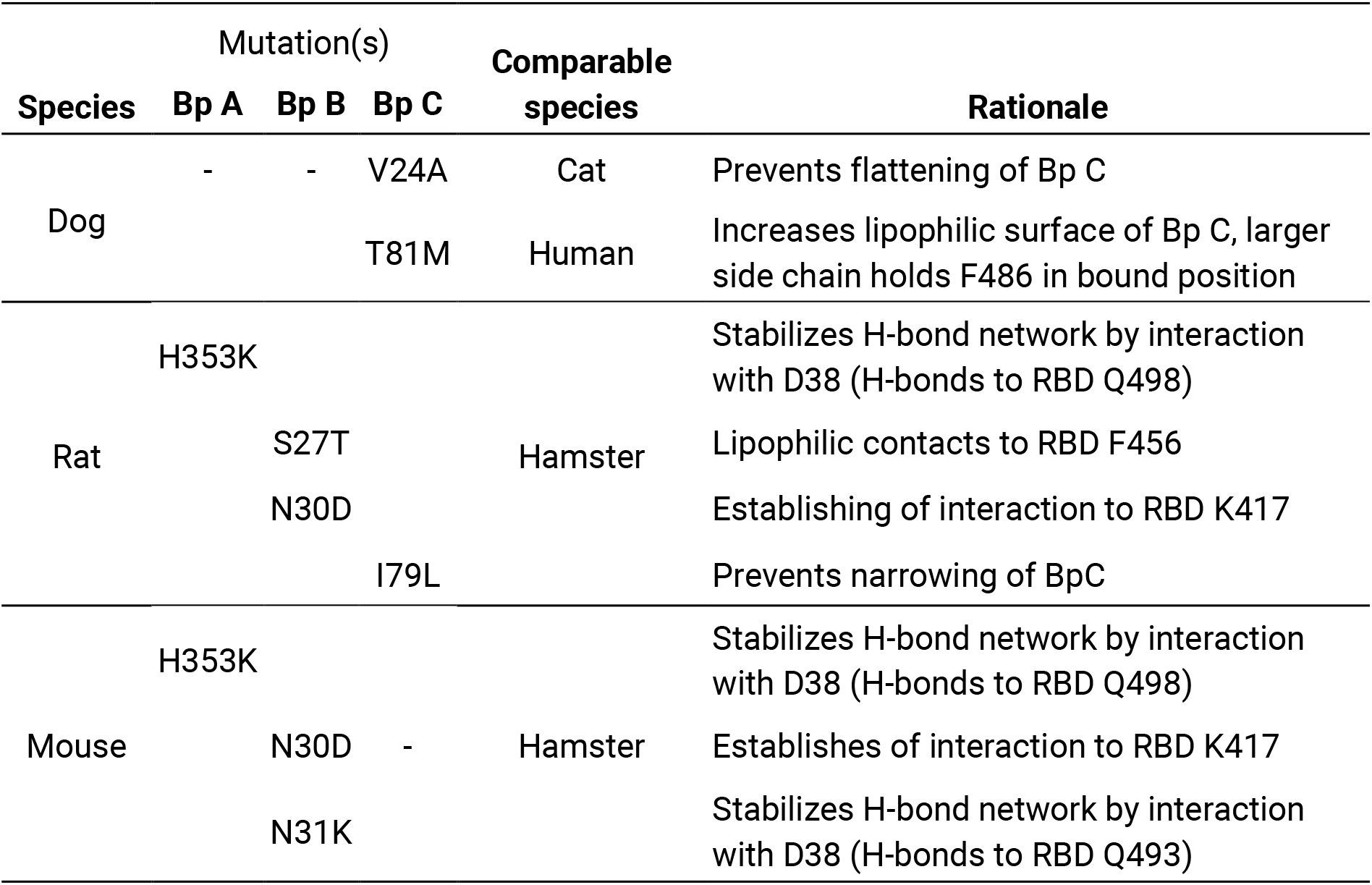
Mutations enhancing SARS-CoV-2 susceptibility suggested by our dynamic models.

In a first step, we mutated residue 24 of canine ACE2 from valine to alanine. This polymorphism should prevent the rotation of Y82 and corresponding of Bp C. The V24A mutant showed the predicted effect (Figure 9). In the 82-78-SDpF torsion diagram of mutated ACE2, the additional peak associated to deformation cannot be observed, which is comparable to feline RBD binding. Moreover, we surmise that mutation T81M in dogs increases susceptibility to SARS-CoV-2 in a fashion that is similar to that conferred by human ACE2.

**Figure 9.**
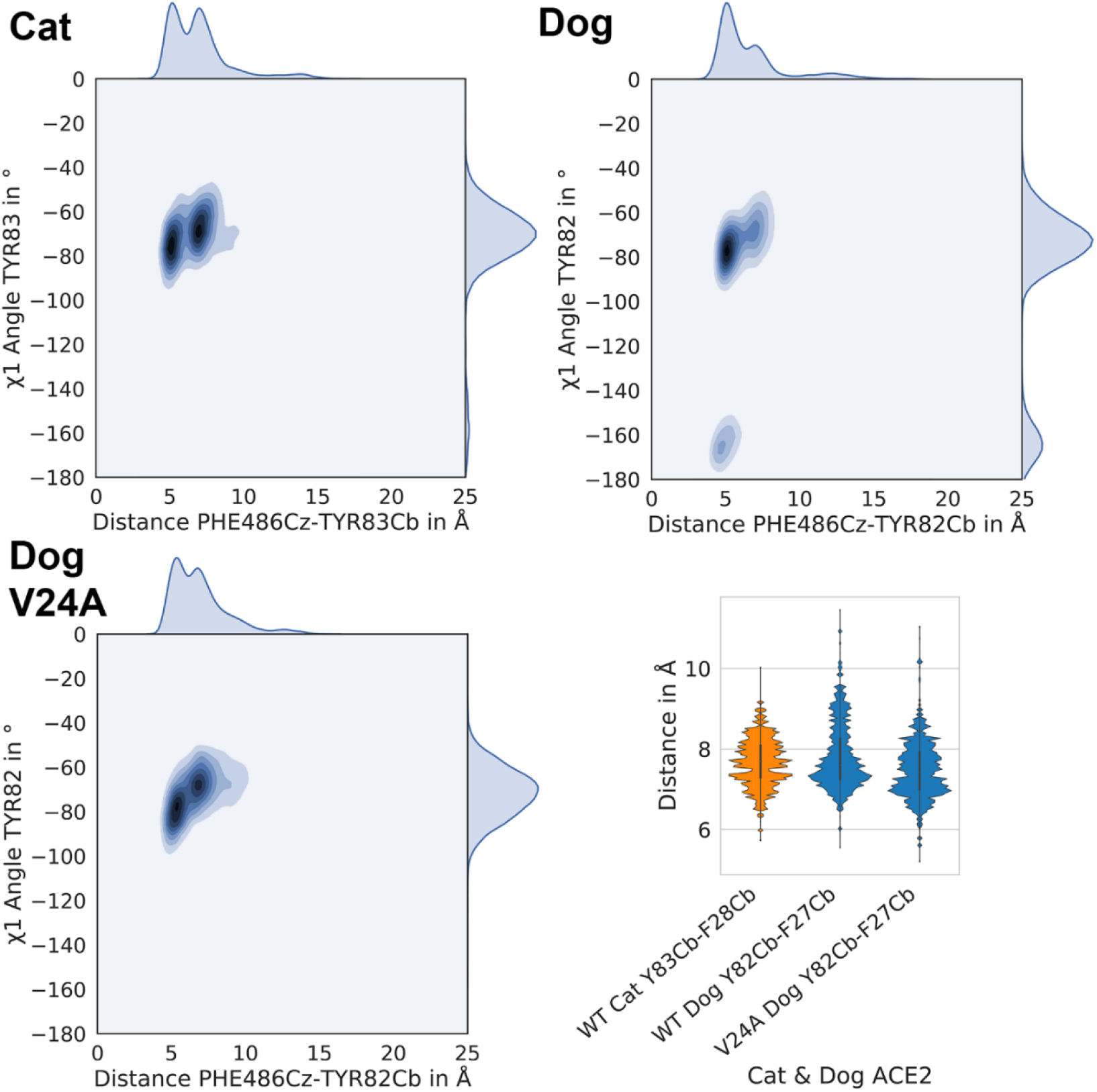
Occupation of binding pocket C indicated by kernel density plots (x-axis: distance F486 Cz - Y82 Cb; y-axis: χ1 angle of central residue 82/83) and Bp C deformation of in dog ACE2 wild type, V24A mutant, and homologous residue 83 in cat (Y83/82 Cb – F28/27 Cb distance)

To test the hypothesis of narrowing the Bp C by branched side chains at position 79, we prepared and simulated an I79L mutant of rat ACE2. This mutation clearly increased Bp C occupancy and showed wide opening of Bp C (Figure 10), which supported our hypothesis.

**Figure 10.**
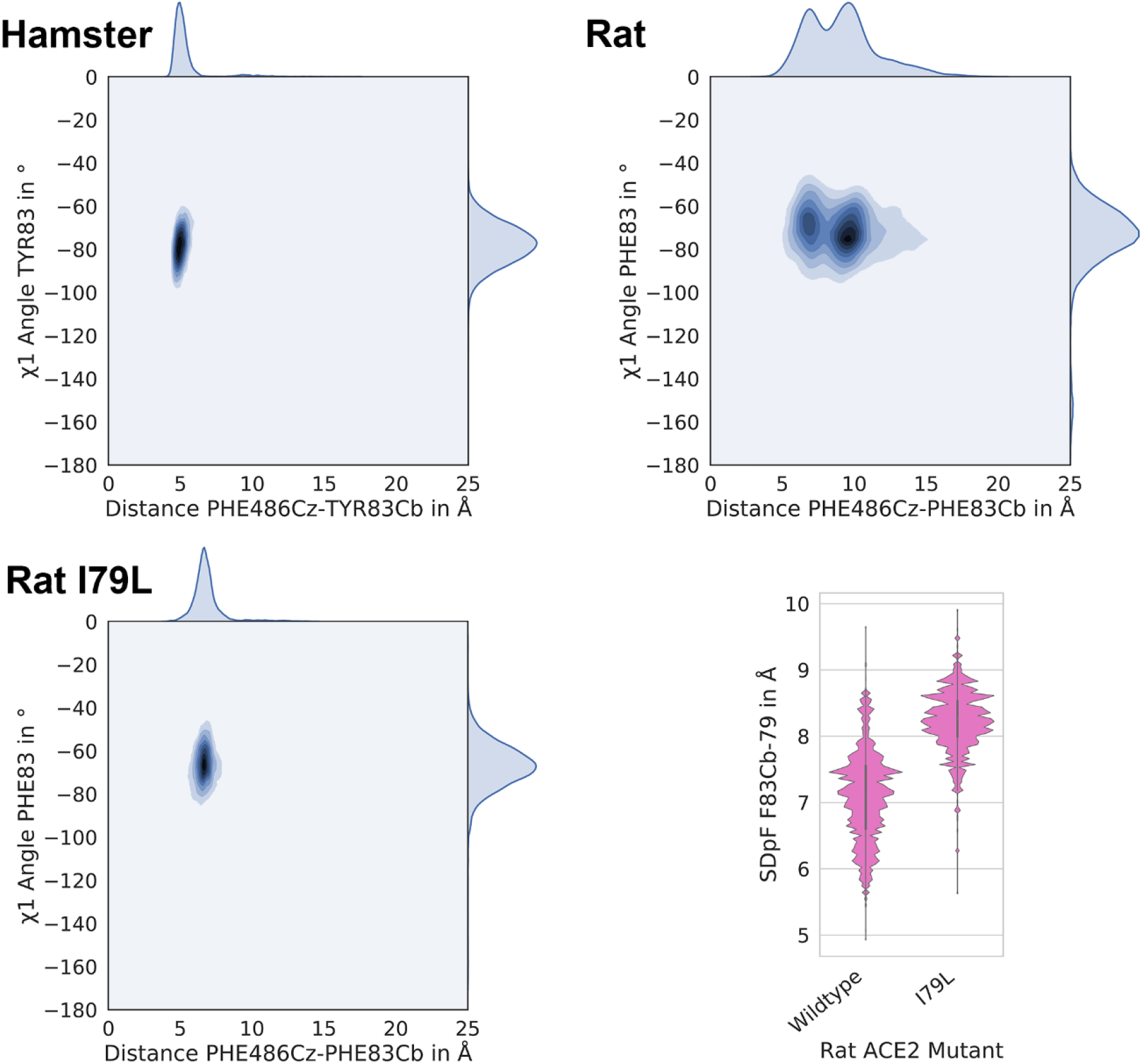
Kernel density plots summarizing the occupation of binding pocket C (x-axis, surrogate parameter: distance F486 Cz - Y83 Cb) and rotamers of central residue 82 (or 83 in cats). The lower right panel shows opening of the Bp C based on the SDpF-83-79 (Shortest Distance per Frame) parameter.

We further suggest to mutate rat and mouse ACE2 in Bp A and B to stabilize RBD binding. In Bp B, mutating positions 30 and 31 from neutral amino acids back to charged ones, as is the case in hamster ACE2, might increase SARS-CoV-2 susceptibility of rat and mouse. Similarly, the mutation H353K in rat and mouse ACE2 Bp A should result in a hydrogen bond network comparable to hamster protein and therefore enhancing the RBD binding.

## Conclusion

We present mechanistic, dynamic models on atomistic level to understand of the ACE2/SARS-CoV-2 interaction in different animal species that might serve as natural reservoirs for SARS-CoV-2 due to frequent contacts with humans. In addition to previous studies^13^, we present extensive molecular dynamics simulations that rationalize RBD binding to ACE2. Based on known susceptibility of animal species to SARS-CoV-2 and the comparison of MD trajectories, we were able to develop models for prediction of RBD binding to ACE2. Hence, we propose gain-of-function mutations for non-susceptible species (dog, rat and mouse) as validation for our model and predict that the red squirrel has a high chance to get infected by SARS-CoV-2.

## Methods

### Homology Modeling and ACE2-Spike Models

Homology and loop models were prepared using MOE 2019. [Chemical Computing Group, Montreal, Canada] The models were constructed using GB/VI scoring^14^ with a maximum of ten main chain models. ACE2 models of cat, dog and ferret were prepared using human ACE2 bound to RBD of SARS-CoV-2 (PDB-ID: 6M0J^5^) as template. ACE2-S complex models were prepared via superposition of receptor homology models on template crystal structure and removing human protein. The catalytic center as well as clashing side chains in binding interface were adjusted using rotamere tool and OPLS-AA force field^15^ in MOE.

### Molecular Dynamics Simulations

All systems (including substrate- and hit-protease complexes) were prepared in MOE 2019 by protonation (at 300 K and pH of 7) and capping protease termini. ACE2-RBD-complex simulations were prepared with Maestro 11.7 [Schrödinger, LLC: New York, USA] and carried out with Desmond 5.5.^16^ All systems were inspected for atom clashes, optimized for H-bonds, filled in 12 Å large padding box with SCP water model^17^, sodium chloride 0.15 M and sodium ions to keep isotonic and electrostatic neutral conditions. The simulations were performed under default minimizing protocol and periodic boundary conditions over 100 ns resulting in 2000 frames each simulation. For visual inspection, all trajectories were wrapped and aligned on backbone Cα atoms of ACE2-RBD-complex and first simulation frame using VMD 1.9.3.^18^

### Trajectory Analysis

Initially, trajectories were analyzed visually to find possible differences in dynamic complexes, such as backbone movements. Further analysis was performed with Python 3.7^19^ using MDAnalysis 0.19.3 for the extraction of distances, angles, and hydrogen bonds from trajectories after an equilibration period of 10 ns (resulting in 1800 complex conformations per replicon). Data processing and transformation was done with pandas 0.25.3.^20^ Plots were created with seaborn 0.9.0.^21^ and matplotlib 3.1.1.^22^

## Notes

### Competing Interest Statement

The authors have declared no competing interest.

## References

1. Dong, E.; Du, H.; Gardner, L., An interactive web-based dashboard to track COVID-19 in real time. The Lancet Infectious Diseases 2020.

2. Zhou, P.; Yang, X. L.; Wang, X. G.; Hu, B.; Zhang, L.; Zhang, W.; Si, H. R.; Zhu, Y.; Li, B.; Huang, C. L.; Chen, H. D.; Chen, J.; Luo, Y.; Guo, H.; Jiang, R. D.; Liu, M. Q.; Chen, Y.; Shen, X. R.; Wang, X.; Zheng, X. S.; Zhao, K.; Chen, Q. J.; Deng, F.; Liu, L. L.; Yan, B.; Zhan, F. X.; Wang, Y. Y.; Xiao, G. F.; Shi, Z. L., A pneumonia outbreak associated with a new coronavirus of probable bat origin. Nature 2020, 579 (7798), 270–273.

3. Gallagher, T. M.; Buchmeier, M. J., Coronavirus spike proteins in viral entry and pathogenesis. Virology 2001, 279 (2), 371–4.

4. Song, W.; Gui, M.; Wang, X.; Xiang, Y., Cryo-EM structure of the SARS coronavirus spike glycoprotein in complex with its host cell receptor ACE2. PLoS Pathog 2018, 14 (8), e1007236.

5. Lan, J.; Ge, J.; Yu, J.; Shan, S.; Zhou, H.; Fan, S.; Zhang, Q.; Shi, X.; Wang, Q.; Zhang, L.; Wang, X., Structure of the SARS-CoV-2 spike receptor-binding domain bound to the ACE2 receptor. Nature 2020.

6. Chen, Y.; Guo, Y.; Pan, Y.; Zhao, Z. J., Structure analysis of the receptor binding of 2019-nCoV. Biochem Biophys Res Commun 2020.

7. Tian, X.; Li, C.; Huang, A.; Xia, S.; Lu, S.; Shi, Z.; Lu, L.; Jiang, S.; Yang, Z.; Wu, Y.; Ying, T., Potent binding of 2019 novel coronavirus spike protein by a SARS coronavirus-specific human monoclonal antibody. Emerg Microbes Infect 2020, 9 (1), 382–385.

8. Towler, P.; Staker, B.; Prasad, S. G.; Menon, S.; Tang, J.; Parsons, T.; Ryan, D.; Fisher, M.; Williams, D.; Dales, N. A.; Patane, M. A.; Pantoliano, M. W., ACE2 X-ray structures reveal a large hinge-bending motion important for inhibitor binding and catalysis. J Biol Chem 2004, 279 (17), 17996–8007.

9. Li, F.; Li, W.; Farzan, M.; Harrison, S. C., Structure of SARS coronavirus spike receptor-binding domain complexed with receptor. Science 2005, 309 (5742), 1864–8.

10. Shi, J.; Wen, Z.; Zhong, G.; Yang, H.; Wang, C.; Huang, B.; Liu, R.; He, X.; Shuai, L.; Sun, Z.; Zhao, Y.; Liu, P.; Liang, L.; Cui, P.; Wang, J.; Zhang, X.; Guan, Y.; Tan, W.; Wu, G.; Chen, H.; Bu, Z., Susceptibility of ferrets, cats, dogs, and other domesticated animals to SARS-coronavirus 2. Science 2020.

11. Cohen, J., Mice, hamsters, ferrets, monkeys. Which lab animals can help defeat the new coronavirus? Science 2020.

12. Bogan, A. A.; Thorn, K. S., Anatomy of hot spots in protein interfaces. J Mol Biol 1998, 280 (1), 1–9.

13. Damas, J.; Hughes, G. M.; Keough, K. C.; Painter, C. A.; Persky, N. S.; Corbo, M.; Hiller, M.; Koepfli, K.-P.; Pfenning, A. R.; Zhao, H.; Genereux, D. P.; Swofford, R.; Pollard, K. S.; Ryder, O. A.; Nweeia, M. T.; Lindblad-Toh, K.; Teeling, E. C.; Karlsson, E. K.; Lewin, H. A., BioRxiv 2020.

14. Labute, P., The generalized Born/volume integral implicit solvent model: estimation of the free energy of hydration using London dispersion instead of atomic surface area. J Comput Chem 2008, 29 (10), 1693–8.

15. Kaminski, G. A.; Friesner, R. A.; Tirado-Rives, J.; Jorgensen, W. L., Evaluation and Reparametrization of the OPLS-AA Force Field for Proteins via Comparison with Accurate Quantum Chemical Calculations on Peptides†. The Journal of Physical Chemistry B 2001, 105 (28), 6474–6487.

16. Bowers, K. J. C., E.; Xu, H.; Dror, R. O.; Eastwood, M. P.; Gregersen, B. A.; Klepeis, J. L.; Kolossvary, I.; Moraes, M. A.; Sacerdoti, F. D.; Salmon, J. K.; Shan, Y.; Shaw, D. E. Scalable Algorithms for Molecular Dynamics Simulations on Commodity Clusters Proceedings of the ACM/IEEE Conference on Supercomputing (SC06), Tampa, Florida, 2006.

17. Toukan, K.; Rahman, A., Molecular-dynamics study of atomic motions in water. Phys Rev B Condens Matter 1985, 31 (5), 2643–2648.

18. Humphrey, W.; Dalke, A.; Schulten, K., VMD: visual molecular dynamics. J Mol Graph 1996, 14 (1), 33–8, 27-8.

19. G. van Rossum, Python tutorial, Technical Report CS-R9526, Centrum voor Wiskunde en Informatica (CWI), Amsterdam, May 1995.

20. McKinney, W., Data structures for statistical computing in python. Proc. of the 9th Python in Science Conf. 2010, 56–61.

21. Waskom, M., mwaskom/seaborn: v0.9.0. https://zenodo.org/record/1313201 2018.

22. Hunter, J. D., Matplotlib: A 2D Graphics Environment. Computing in Science & Engineering 2007, 9 (3), 90–95.

